# Inhibition of PIM kinase in tumor associated macrophages suppresses inflammasome activation and sensitizes prostate cancer to immunotherapy

**DOI:** 10.1101/2024.10.21.618756

**Authors:** Amber N. Clements, Andrea L. Casillas, Caitlyn E. Flores, Hope Liou, Rachel K. Toth, Shailender S. Chauhan, Kai Sutterby, Sachin Kumar Deshmukh, Sharon Wu, Joanne Xiu, Alex Farrell, Milan Radovich, Chadi Nabhan, Elisabeth I. Heath, Rana R. McKay, Noor Subah, Sara Centuori, Travis J. Wheeler, Anne E. Cress, Gregory C. Rogers, Justin E. Wilson, Alejandro Recio-Boiles, Noel A. Warfel

**Author notes:** The authors declare no conflict of interest.

## Abstract

Immunotherapy has changed the treatment paradigm for many types of cancer, but immune checkpoint inhibitors (ICIs) have not shown benefit in prostate cancer (PCa). Chronic inflammation contributes to the immunosuppressive prostate tumor microenvironment (TME) and is associated with poor response to ICIs. The primary source of inflammatory cytokine production is the inflammasome. Here, we identify PIM kinases as important regulators of inflammasome activation in tumor associated macrophages (TAMs). Analysis of clinical data from a cohort of treatment naïve, hormone responsive PCa patients revealed that tumors from patients with high PIM1/2/3 display an immunosuppressive TME characterized by high inflammation (IL-1β and TNFα) and a high density of repressive immune cells, most notably TAMs. Strikingly, macrophage-specific knockout of PIM reduced tumor growth in syngeneic models of prostate cancer. Transcriptional analyses indicate that eliminating PIM from macrophages enhanced the adaptive immune response and increased cytotoxic immune cells. Combined treatment with PIM inhibitors and ICIs synergistically reduced tumor growth. Immune profiling revealed that PIM inhibitors sensitized PCa tumors to ICIs by increasing tumor suppressive TAMs and increasing the activation of cytotoxic T cells. Collectively, our data implicate macrophage PIM as a driver of inflammation that limits the potency of ICIs and provides preclinical evidence that PIM inhibitors are an effective strategy to improve the efficacy of immunotherapy in prostate cancer.

## Introduction

Prostate tumors are commonly termed immunologically ‘cold’ due to a lack of cytotoxic T cell infiltration and high numbers of immunosuppressive immune cells, including myeloid derived suppressor cells (MDSCs), regulatory T cells (Tregs), and tumor associated macrophages (TAMs). High TAM density is prevalent in PCa and is correlated with worse recurrence free survival following hormone therapy (1,2). TAMs inhibit anti-tumor responses through direct and indirect mechanisms. One mechanism by which TAMs promote tumor growth is by inducing chronic inflammation, which activates immune checkpoint pathways and increases the secretion of inhibitory cytokines that suppress the immune response (3). Due to their importance in prostate cancer progression and therapeutic resistance, eliminating or reprogramming TAMs has been a long-sought after goal for prostate cancer therapy (4).

Chronic inflammation is an established mechanism of resistance to immune checkpoint inhibitors (ICIs) (5–7). Upregulation of inflammatory cytokines and chemokines in tumor cells and surrounding immune cells (namely, TAMs) is a hallmark of cancer. Macrophages express NLRP3, a cytosolic immune sensor that oligomerizes to form the inflammasome (8). Indicative of its pro-tumorigenic functions, NLRP3 expression is upregulated in human PCa and enhances cell growth, survival, and migration of prostate cancer cells (9). Moreover, serum levels of the proinflammatory cytokines that are released by inflammasomes, including IL-1β and IL-18, are significantly higher in patients with locally advanced PCa compared to healthy controls, suggesting that activation of the inflammasome is associated with PCa progression (10). High intratumoral levels of IL-1β is an established driver of PCa growth, and high IL-1β is significantly associated with biochemical recurrence in PCa patients (11). In the context of the tumor-immune microenvironment, high levels of IL-1β is correlated with increased numbers of immunosuppressive immune cells and worse response to ICIs in PCa (12–14). These studies demonstrate that aberrant activation of inflammasomes during prostate tumorigenesis is a key factor contributing to inflammation and the immunosuppressive prostate TME.

The Proviral Integration site for Moloney murine leukemia virus (PIM) kinases are a family of oncogenic Ser/Thr kinases (PIM1, PIM2, PIM3) whose levels are elevated in high-grade prostatic intraepithelial neoplasia (HGPIN) relative to normal tissue and further elevated in mCRPC (15,16). PIM regulates signaling pathways that promote cell proliferation and survival in cancer cells, and recent studies have linked increased PIM expression to immune evasion through tumor-intrinsic mechanisms (17–19). However, the role of PIM in tumor-associated immune cells and its impact on response to immune checkpoint inhibitors (ICIs) is not well known (19–22). The PIM isoforms are encoded by genes located on different chromosomes. At the amino acid level, the PIM isoforms have high sequence similarity. Specifically, PIM1 and PIM2 have 61% sequence homology, while PIM1 and PIM3 are 71% identical (23). Growing evidence indicates that PIM is involved in controlling inflammation in normal and tumor tissues. Transgenic mouse models overexpressing PIM1 or PIM2 in hormone-dependent tissues show increased inflammatory immune cell infiltration (24,25), and high PIM1 is correlated with inflammatory features in human breast and prostate tumors (26).

In this study, we demonstrated expression of PIM in TAMs is particularly important in creating an immunosuppressive TME in PCa. Clinical data show that high PIM expression is correlated with inflammation, excessive TAMs, and lack of cytotoxic immune cells. Using genetic and pharmacological approaches, we show that macrophage-specific loss of PIM suppressed tumor growth in syngeneic co-injection models *in vivo*. Transcriptional analysis of tumor tissue and biochemical studies using bone marrow derived macrophages (BMDM) indicate that PIM increases inflammasome activation and the release of inflammatory cytokines. Combining PIM inhibitors with ICIs produced synergistic anti-tumor effects, characterized by reduced repressive immune cells and increased cytotoxic T cells. Collectively, our data implicate PIM as a limiting factor for the potency of ICIs and provide preclinical evidence to support further testing of PIM inhibitors as a strategy to realize the benefit of immunotherapy in prostate cancer.

## Materials and Methods

### Cell culture

MyC-CaP (ATCC CRL-3255) prostate cancer cells were cultured in DMEM (Cat# 10-017-CV; Corning) supplemented with 10% Fetal Bovine Serum (Omega Scientific) in a humidified atmosphere of 5% CO2 at 37°C. Bone marrow cells were isolated from the femurs and tibias of WT FVB/J mice and differentiated in culture for 6-7 days in RPMI 1640 (Corning) supplemented with 10% heat inactivated FBS, 2mM L-glutamine, 1% penicillin/streptomycin, and with 15% CMG14-12 conditioned media. Differentiated bone marrow derived macrophages (BMDMs) were collected and plated for downstream analysis.

### Mice

Pim single KO and Pim1-/-Pim2-/-Pim3-/-triple KO (TKO) mice were generated by Mikkers, et al. (Mikkers, Nawijn et al. 2004) and were kindly provided by Andrew S. Kraft. FVB/J mice used for all *in vivo* experiments were purchased from the Jackson Laboratory. All animal studies were approved by the Institutional Animal Care and Use Committee at the University of Arizona.

### BMDM polarization and inflammasome activation

Bone marrow derived macrophages were treated with IFNγ (50ng/ml) (PeproTech) and lipopolysaccharide (LPS) (20ng/ml) (Invivogen) or IL-4 (20ng/ml) (PeproTech) for 24 hours to skew towards M1 and M2 macrophages, respectively. For inflammasome activation, BMDMs were treated with LPS (500ng/ml) (Invivogen) for 4 hours, followed by stimulation with ATP (2mM) (Invivogen) for 30 minutes. To inhibit PIM activity, BMDMs were treated overnight (16 hours) with PIM447 (3μM) (MedChemExpress) prior to inflammasome activation. Supernatant was collected and mouse IL-1β levels were measured using an ELISA kit (R&D Systems, Minneapolis, MN, USA, DY401). Protein lysates were also collected for immunoblotting analysis.

### Patient data analysis

44 patients of treatment-naive metastatic hormone-sensitive PC were analyzed by DNA next-generation sequencing (592-gene, NextSeq; WES, NovaSeq) and Whole Transcriptome Sequencing (WTS; NovaSeq) (Caris Life Sciences, Phoenix, AZ). PC with PIM1/2/3-high(H) N28 and -low(L) N16 expressions were classified by top and bottom quartile, respectively. Androgen receptor (AR) signature and Neuroendocrine Prostate Cancer (NEPC) score were calculated based on the expression level of previously defined gene signatures (Hieronymus et al. 2006, Beltran et al. 2016). T cell inflamed score and IFNγ score were calculated based on previously defined parameters (Bao et al 2020, Cristescu et al 2018). Pathway enrichment was determined by GSEA (Broad Inst). Immune cell fractions were calculated by deconvolution of WTS using Quantiseq. Statistical significance was determined by chi-square and Mann-Whitney U (p<0.05) and adjusted for multiple comparisons (p<0.05). Continuous and categorical variables were evaluated using t-tests or Kruskal-Wallis Rank sum tests and Chi-square or Fisher’s exact tests, respectively. All analyses were conducted using Stata 18.

### Immunoblotting

Cell lysates were prepared with RIPA buffer (1X PBS, 150mM NaCl, 1% Nonidet P40, 0.5% NaCl, 0.5% sodium deocycholate, 0.1% SDS, 25mM Tris) and protease inhibitor cocktail (Fisher Scientific). Total protein concentration was measured using BCA Protein Assay Kit (Cat 23225: Pierce, Rockford). Equal protein concentrations were loaded and separated on SDS-PAGE gels and blotted onto 0.45μm PVDF transfer membranes (Cat: 88518, Thermo Scientific). Membranes were blocked in 5% non-fat milk in Tris-buffered saline with tween (TBST) for one hour. Primary antibodies were diluted in 5% Bovine Serum Albumin (BSA) and incubated overnight at 4°C. The following primary antibodies were used: anti-phospho-STAT1 (Tyr701)(Cell signaling; 1:1,000), anti-total STAT1 (Cell Signaling; 1:1000), anti-phospho-STAT6 (Tyr641) (Cell Signaling; 1:1000), anti-total STAT6 (Cell Signaling; 1:1000), anti-caspase-1 (p20) (mouse) (AdipoGen; 1:2000), anti-NLRP3 (D4D8T) (Cell Signaling), anti-Gasdermin D (E9S1X) (Cell Signaling;1:1000), anti-Nf-κB p65 (L8F6) (Cell Signaling; 1:1000), anti-Phospho-NF-kB p65 (Ser536) (Cell Signaling; 1:1000), anti-Actin (BD Biosciences;1:5000). Following washing in TBST, membranes were incubated with secondary antibody (anti-Rabbit IgG HRP-linked, Cell Signaling or anti-mouse HRP-linked, Genesee Scientific; 1:5000) for 1 hour. Membranes were incubated with ECL Substrate (Bio-Rad) or SuperSignal West Femto Chemiluminescent Substrate (Fisher Scientific) and imaged using the Syngene G:Box.

### qRT-PCR Analysis

Total RNA was isolated from cell lysates using the Quick-RNA MiniPrep Kit (Zymo Research; Cat# R0155), and cDNA was synthesized from each sample using qScript™ cDNA SuperMix (QuantaBio). qRT-PCR reactions were performed with equal amounts of starting material (1000ng RNA) using qPCRBIO SyGreen supermix (Genesee Scientific), according to manufacturer’s instructions.

### Myc-CaP co-injection *in vivo*

To deplete resident macrophages, male WT FVB/J mice (Jackson Laboratory) were injected retro-orbitally with clodronate liposomes (Liposoma BV). The following day, a mixture of MyC-Cap and WT, PIM1 knockout, or triple knockout (TKO) BMDMs were injected into the right and left flanks of the FVB mice at a ratio of 2:1. Tumor volume was monitored over time by caliper measurement and calculated as (length x width2)/2. At the end of the experiment, tumors were fixed, embedded in paraffin, and sectioned for staining with hematoxylin and eosin (H&E) or antibodies specific for Ki67, cleaved caspase-3, F4/80 and CD8. Independent investigators blinded to the identity of the samples scored positive staining using QuPath image analysis. Total RNA was isolated from paraffin embedded tumor tissue using deparaffinization solution (Qiagen;Cat#19093) and the RNeasy FFPE Kit (Qiagen;Cat#73504) according to the manufacturer’s instructions. All samples were prepared in nuclease-free water. The RNA samples were submitted to the Arizona Genetics Core to run the NanoString nCounter® mouse immunology panel (NS_Immunology_Mm_C2269). Data analysis and graphs were generated using the nSolver advanced analysis software.

### *In vivo* combination treatment study

MyC-Cap cells (1×10^6^) were injected into the right and left flanks of male WT FVB mice (Jackson Labtoratory). Once the tumors reached a volume of approximately 100 mm3, the mice were randomized for treatment with vehicle and isotype control, PIM447 (30 mg/kg/day by p.o.), Anti-PD-1 Ab (200ug/mouse, i.p. three times a week), or combination of PIM447 and Anti-PD-1 (n=4 mice/group). Tumor volumes were measured over time by caliper measurement and calculated as (length x width2)/2. The mice were euthanized when the tumor measured greater than 2000 mm^3^. The tumors were harvested from the mice and half the tumors from each group were fixed, embedded in paraffin, and sectioned for staining with hematoxylin and eosin (H&E) or antibodies specific for Ki67, cleaved caspase-3, and F4/80. The remaining tumors were minced into small pieces. To form a single cell suspension, the tumors were incubated at 37°C for 30 minutes in RPMI supplemented with 2% FBS, 1mg/ml Collagenase Type 1 (EMD Millipore),. 1 mg/ml DNase, and MgCl. The remaining tissue was gently grinded through a 100 μm cell strainer. Single-cell suspensions were generated for downstream flow cytometry analysis.

### Spectral flow cytometry

Single-cell suspensions were blocked with anti-mouse CD16/32 antibody (TruStain FcXTM PLUS, Cat# 156603, Biolegend) for 10 minutes and incubated with antibodies from the Cytek 17 color mouse immune panel (RC-00152). Intracellular staining was performed after extracellular staining using the True-Nuclear Transcription Factor Staining Buffer Kit (Cat# 424401). The cells were fixed, and all samples were run on the Cytek Aurora full spectrum flow cytometer equipped with five lasers (355, 405, 488, 561 and 640 nm excitation wavelengths), three scattering channels, and detection up to 64 fluorescence channels, housed in the Flow Cytometry and Immune Monitoring Shared Resource (FCIMSR) located in the University of Arizona Cancer Center. Data was analyzed using SpectroFlo (Cytek) and FlowJo software. All antibodies used for the spectral flow cytometry analysis are listed in the Supplementary Table 1.

### Immune profiling analysis

FlowSOM Clustering was performed on spectral flow cytometry data using the OMIQ platform. Thirty individual immune cell populations clusters were generated and data was visualized as UMAP plots. The clusters were tested for significance between each group. Criteria for a significant cluster was adjusted P-value false discovery rate (FDR) <= 0.05. The edgeR method used to test for differential abundance between groups and volcano plots were created showing statistical significance (FDR) on the Y axis versus magnitude of change (Fold change) on the x axis. A positive fold change means upregulated, and a negative means downregulated. Boxplots were generated for significant clusters among groups based on FDR<=0.05.

### Statistical analysis

Statistical analysis and graphical representation of the data were performed in GraphPad Prism. A minimum number of three replicates were performed for all in vitro experiments. One way ANOVA and unpaired t tests were performed unless stated otherwise in the figure legends. Statistical significance is denoted in figures as (*p≤.05,**p≤.01). Figure legends contain information regarding n values and p values. Data are presented as mean ± SEM.

## Results

### High PIM expression correlates with increased inflammation and immunosuppressive cells in human prostate tumors

A major reason that ICIs have not been effective in prostate cancer is that the prostate TME is typically made up of immune cells that suppress the anti-tumor response. To determine whether increased PIM kinase levels contribute to the immunosuppressive nature of PCa, we analyzed whole transcriptomic data collected from the tumors of 44 patients with treatment-naive metastatic hormone-sensitive PCa collected at the University of Arizona Cancer Center (Fig. 1A). Tumors were classified based on PIM1/2/3 expression, and tumors in the top (*PIM*-high) and bottom quartile (*PIM*-low) were compared. Consistent with our previous work showing that PIM1 increases activation of hypoxia-inducible factor-1 (HIF-1), hypoxia-related gene expression was significantly increased in *PIM*-high tumors compared to *PIM*-low tumors (Fig. S1A). (27) High PIM expression was more common in later stage disease (82% *PIM*-high vs. 57% *PIM*-low in patients with Gleason 3+4) and was associated with higher median PSA (66.0 *PIM-*high vs. 24.0 *PIM*-low) (Table 1). *PIM*-high tumors also correlated with increases in clinical metrics of prostate cancer progression, including PSA and androgen receptor (AR) expression, as well as increased MAPK pathway activity (Fig. S1B-E). Notably, *PIM*-high tumors also showed highly elevated makers of chronic inflammation: T cell inflamed score and proinflammatory cytokines (i.e., *IL-1β*, *TNF*, *IFNγ*, and *TNFSF13*) (Fig. 1B, C). These data validate that PIM expression is correlated with markers of disease progression in this patient cohort.

**Figure 1.**
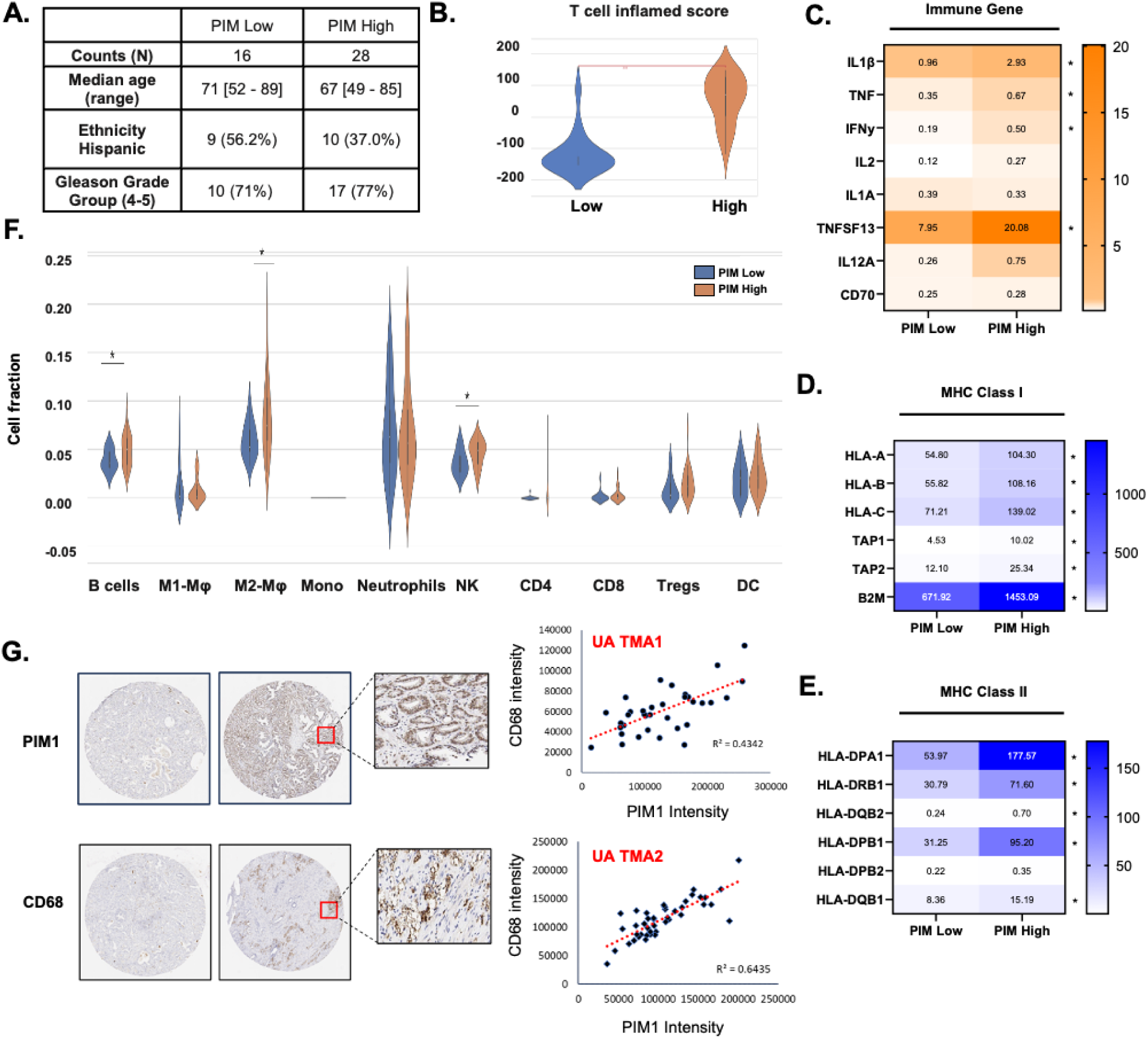
PIM kinase expression correlates with increased inflammation and immune cell infiltration in human prostate cancer. A) 44 treatment-naive metastatic hormone-sensitive PCa pts were analyzed by DNA 592-gene, NextSeq; WES & WTS; NovaSeq. Patient tumors were classified as *PIM*-high and *PIM*-low based on the top and bottom quartile of PIM expression. Violin plots and heatmaps comparing B) T cell inflamed score, C) Immune gene expression, D) MHC class I genes, and E) MHC class II genes. F) Cell fraction of total immune cells in *PIM*-high and *PIM*-low tumors. G) Tissue Microarrays (TMAs; Gleason 3+ to 3+4+7) were stained for CD68 and PIM1 and relative expression in each core is plotted. **p<.01, *p<.05.

**Table 1.** Clinical characteristic of PCa patient cohort.

|  | **PIM-high** | **PIM-low** | **p-value** |
| --- | --- | --- | --- |
| **Counts** | 28 | 16 |  |
| **Mean age (SD)** | 66.15 (10.40) | 69.19 (11.14) | 0.37 |
| **Median age (IQR)** | 65 [58-74] | 69.5 [62.5-76.5] | 0.48 |
| **Ethnicity Hispanic** | 10 (37.0%) | 9 (56.2%) | 0.22 |
| **Gleason Grade Group (4-5)** | 17 (77%) | 10 (71%) | 0.69 |
| **Karnofsky ≥ 90** | 17 (63%) | 5 (31%) | 0.04 |
| **Tumor Stage** |  |  | 0.36 |
| **1** | 0 ( 0%) | 2 (14 %) |  |
| **2** | 3 (18%) | 4 ( 29 %) |  |
| **3** | 8 (47%) | 3 ( 21%) |  |
| **4** | 6 (35%) | 5 (36%) |  |
| **Node Positive** | 16 (76%) | 11 (69%) | 0.72 |
| **Metastasis** |  |  | 0.44 |
| **M0** | 2 ( 8%) | 3 (19%) |  |
| **M1a/M1b** | 14 (54%) | 9 (56%) |  |
| **M1c** | 10 (38%) | 4 (25%) |  |
| **Median PSA [Range]** | 66.0 [14.0-450.7] | 24.0 [6.6-95.0] | 0.14 |

Next, we analyzed how PIM levels influenced immune populations in patient tumors. First, we assessed markers of antigen presentation, which is critical for eliciting effective antitumor responses. *PIM*-high tumors showed higher expression of both human leukocyte antigen (HLA) class I and class II molecules compared to *PIM*-low tumors (Fig. 1D, E). Isolating specific immune populations revealed that *PIM-*high tumors correlated with increased numbers of B cells, natural killer cells, and M2 macrophages (Fig. 1F). Because macrophages are the most prevalent immune cell in prostate cancer, we sought to confirm their association with PIM1 levels in human tumors. Two prostate cancer tissue microarrays (TMAs) containing 30 samples obtained from diagnostic cores of radical prostatectomies of patients prior to treatment (Gleason score 6-9) were stained for PIM1 and CD68 (macrophage marker), and immunohistochemical staining was quantified based on the intensity and area of positive staining. In both TMAs, there was a significant correlation between high PIM1 and increased macrophage density (Fig. 1G). Analysis of publicly available data from The Cancer Genome Atlas (TCGA) prostate adenocarcinoma (PRAD) cohorts further validated that *PIM1* expression correlated with *CD68*, as well as *CD86* (M1 macrophage marker) and *Mrc1* (M2 macrophage marker) in prostate cancer patients (Fig. S1I). Parallel analyses to compare high vs. low expression of individual PIM isoforms (PIM1, PIM2, and PIM3) indicate that they are largely overlapping with immune cell populations, but PIM1 and PIM2 are significantly correlated with macrophage markers, whereas PIM3 was not (Fig S1, S2). Together, these data show that increased PIM expression correlates with inflammation and higher TAM density.

### PIM Expression in Macrophages Promotes Tumor growth

Macrophages are a primary source of inflammatory cytokine production in solid tumors. Due to the strong association between PIM1 levels and macrophages in patient samples, we asked whether macrophage-specific PIM1 promotes inflammation and suppresses immune responses. To test this, we designed an *in vivo* co-injection experiment. All mice were first treated with clodronate liposomes to deplete resident macrophages. Bone marrow was isolated from wild type, PIM1 knockout (KO) and PIM triple knockout (TKO; lacking all 3 isoforms) mice. Monocytes were collected and differentiated into macrophages following established protocols (28). Then, macrophages (1 x 10^5^ cells) and MyC-CaP cells (1 x 10^6^ cells) were mixed, injected into the flanks of wild type FVB mice, and tumor volume was measured over time (Fig. 2A). The addition of WT macrophages to MyC-CaP increased tumor growth compared to MyC-Cap alone (Fig 2B). Tumors containing PIM1 KO macrophages grew at a similar rate as MyC-Cap only tumors, suggesting that losing PIM1 eliminates the growth advantage provided by macrophages (Fig. 2B). Surprisingly, tumors containing TKO macrophages were significantly smaller than MyC-CaP alone, indicating that loss of PIM in macrophages is sufficient to suppress tumor growth (Fig 2B). At the end of the study, tumors were collected for immunohistochemical analysis. As expected, co-injection tumors compared contained more macrophages (F4/80^+^) than MyC-CaP tumors alone (Fig 2C, D). Tumors from the TKO co-injection group showed decreased cell proliferation (Ki67) and increased cell death (cleaved-caspase 3) compared to MyC-CaP controls. Notably, the number of cytotoxic CD8+ T cells was significantly higher in TKO co-injection tumors compared to WT or control tumors (Fig. 2C, D). To determine how loss of macrophage PIM affected immune signaling, RNA was isolated from each tumor for NanoString nCounter analysis using a mouse-specific immunology panel. Immune signaling genes were highly upregulated in TKO co-injected tumors compared to all other groups (Fig. 2E). Pathway analysis revealed that genes related to apoptosis, lymphocyte trafficking, adaptive immune signaling, and lymphocyte activation were significantly upregulated in TKO co-injected tumors (Fig. 2F). Using immune cell specific gene-signatures, we observed more CD45+ cells in TKO co-injection tumors, particularly increased neutrophils and cytotoxic immune cells (Fig. S3A). In addition, TKO co-injected tumors showed increased expression of cytokines related to the innate immune response, the most significant being *CCL24* (7.48 log2 fold change), a chemoattractant for resting T lymphocytes, eosinophils, and neutrophils (Fig. S3B, S3C) (29). Collectively, these data indicate that PIM is essential for the pro-tumorigenic effect of TAMs, and loss of PIM in macrophages suppresses the tumor immune response.

**Figure 2.**
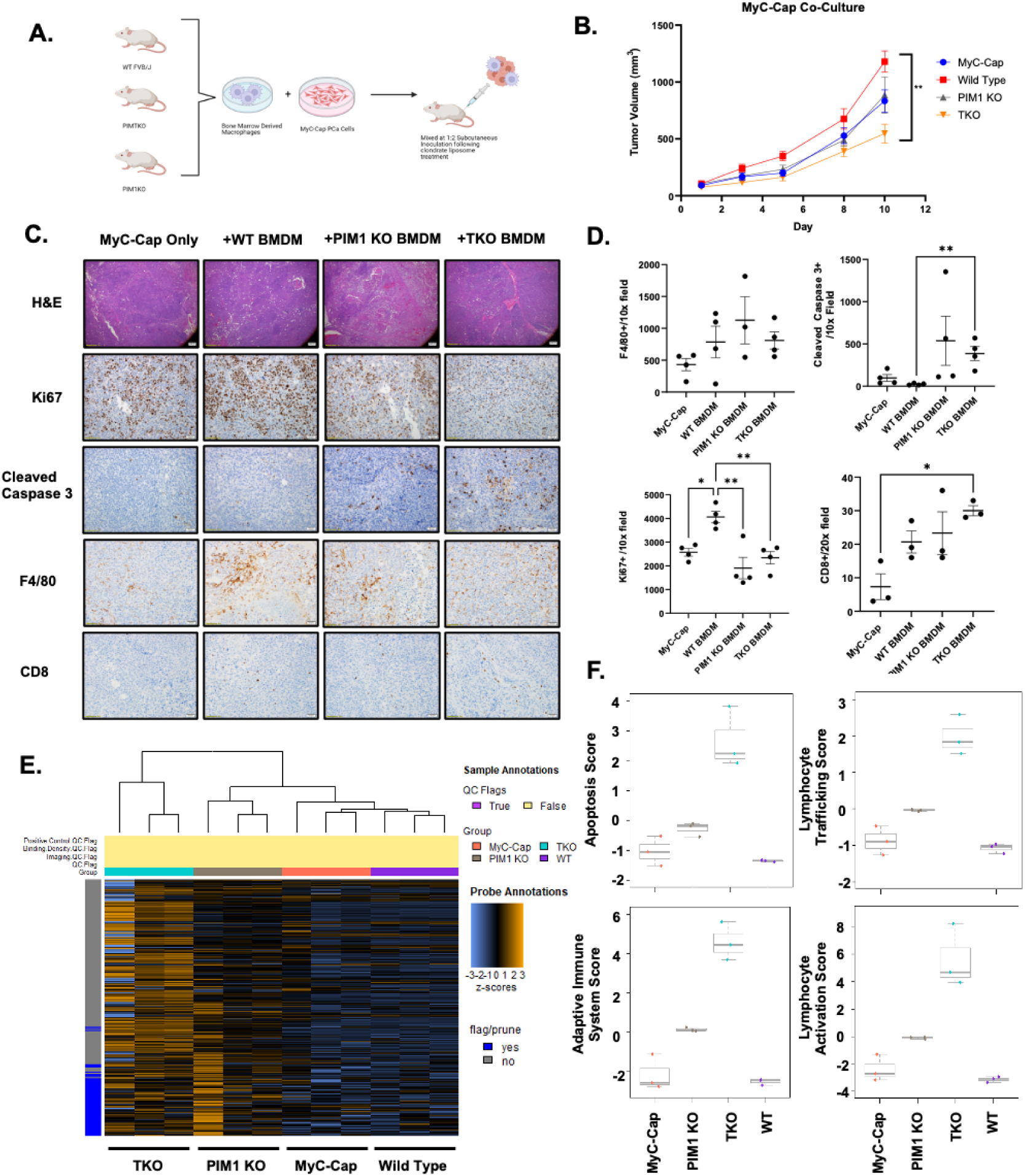
Loss of PIM expression in macrophages activates the immune response and suppresses tumor growth. A) Schematic illustration of the tumor co-injection model. B) BMDMs generated from wild type (WT), PIM1 knockout (PIM1 KO), and triple knockout (TKO) mice (5×10^5^) were co-injected with MyC-CaP (1×10^6^) prostate cancer and tumor volume was measured over time. C and D)) Staining and quantification of Ki67, Cleaved Caspase 3, F4/80, and CD8 immunostaining in tumor from each cohort. E) Heat map showing gene expression changes detected by nCounter analysis in co-injected tumors, F) Apoptosis, lymphocyte trafficking, adaptive immune system, and lymphocyte activation scores (n=3/group); Box plots represent the 25^th^, 50^th^ (median), and 75^th^ percentiles. (*p<.05,**p<.01).

### PIM inhibition suppresses inflammasome activation in bone marrow derived macrophages

Tumor associated macrophages can have positive and negative effects on tumor growth depending on their polarization state, which is generally described as being M1 (anti-tumor) or M2 (pro-tumor). Thus, we reasoned that PIM could alter inflammatory signaling by shifting macrophage polarization state. To test this, WT and TKO macrophages were skewed toward M1 using IFNγ (50 ng/ml) and lipopolysaccharide (LPS; 20 ng/ml for 24 hours) or toward M2 by treatment with IL-4 (20 ng/ml for 24 hours) and the extent of polarization was determine by assessing the phosphorylation state of STAT1 (Y701) and STAT6 (Y641), which are responsible for the transcription of M1 and M2 target genes, respectively(30). Following M1 stimulation, TKO macrophages displayed 2-fold higher phosphorylation of STAT1 (Y701) compared to WT macrophages, suggesting that loss of PIM leads to a more anti-tumor, M1 phenotype (Fig. 3A). In contrast, phosphorylation of STAT6 (Y641) was similar in WT and TKO macrophages skewed towards M2 (Fig. 3B). Next, we next assessed polarization in WT BMDMs after treatment with PIM447, a pan-PIM kinase inhibitor. Like genetic loss of PIM, pharmacological inhibition significantly increased phosphorylation of STAT1 (Y701) but did not alter phosphorylation of STAT6 (Y641) when stimulated with IL-4. Interestingly, phosphorylation of STAT6 was increased in unstimulated macrophages (Fig 3C), suggesting that PIM inhibitors would direct naïve monocytes toward an M2 state. Overall, these data indicate that blocking PIM shifts TAMs towards and M1 phenotype.

**Figure 3.**
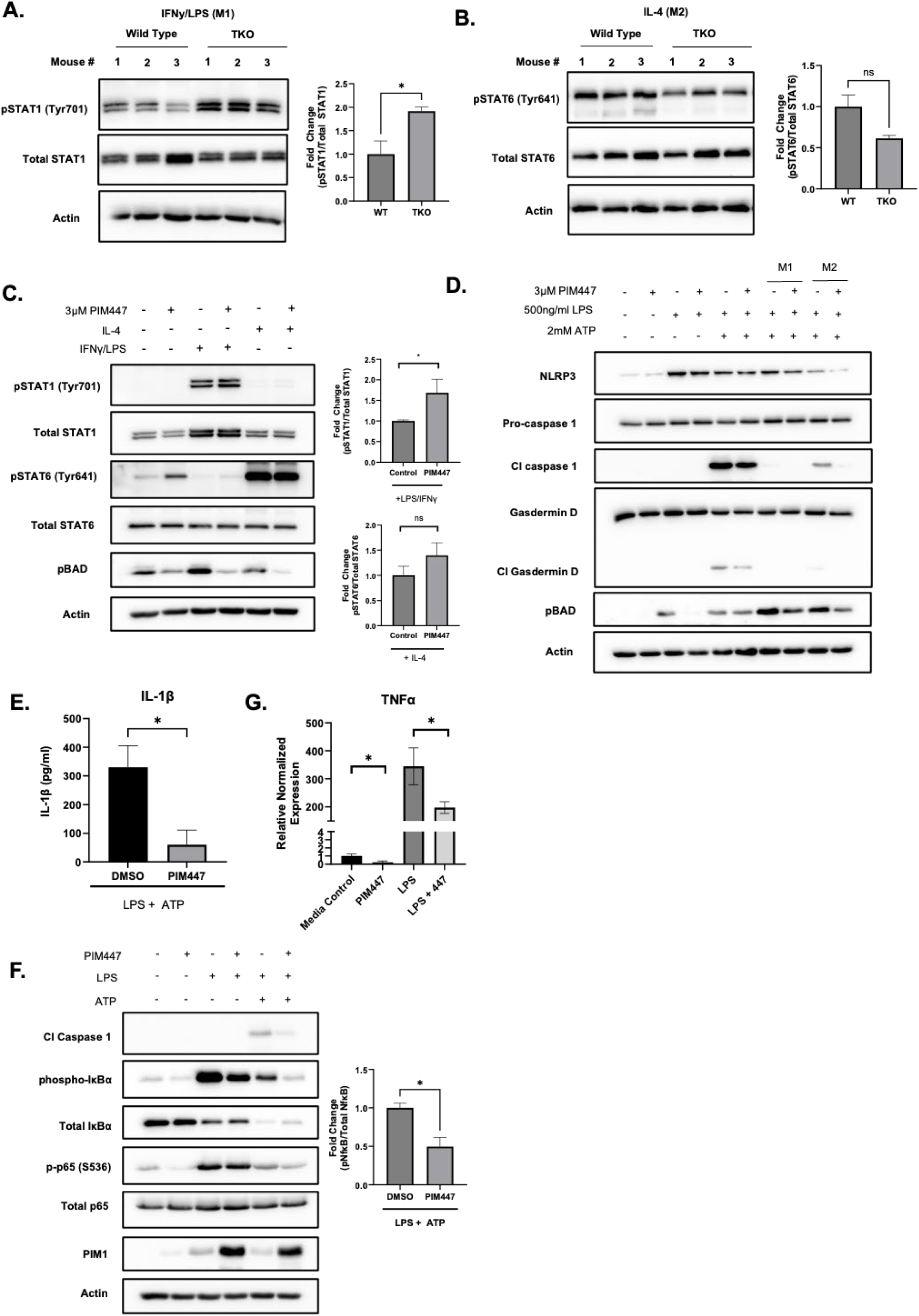
Genetic or pharmacological inhibition of PIM suppresses inflammasome activation. A) Wild type (WT) and PIM triple knockout (TKO) BMDMs were stimulated with IFNγ (500 ng/ml) and LPS) (20ng/ml) for 24 hours or B) with IL-4 (20 ng/ml) for 24 hours, and lysates were collected for immunoblotting C) WT BMDMs were treated with PIM447 (3μM) and stimulated with IFNγ/LPS (M1) or IL-4 (M2). Lysates were analyzed by western blotting. D) WT BMDMs were pre-treated with PIM447 for 16 hours, followed by stimulation with LPS for 4 hours and ATP for 30 min to induce inflammasome activation. Lysates were analyzed by western blotting. E) Supernatant was collected from each cell condition and secreted IL-1β was quantified by ELISA. F) WT BMDM were pre-treated with PIM447, followed by LPS/ATP and immunoblotting was used to assess NF-κB proteins. G) mRNA analysis of TNFα.*p<0.05 Error bars = SEM.

The association between PIM and pro-inflammatory molecules that are commonly released from macrophages led us to test whether PIM can regulate the inflammasome. The inflammasome is an innate immune receptor that is responsible for cleaving IL-1β and other factors into their bioactive form. To stimulate inflammasome activation, WT BMDMs were treated with LPS (500 ng/ml) for 3 hours and ATP (2 mM) for 30 minutes, which caused a robust increase of cleavage of inflammasome substrates, caspase 1 and gasdermin D. However, pre-treatment with PIM447 (3 μM) for 16 hours significantly blunted inflammasome activation (Fig 3D). ELISA assays confirmed that PIM447 treatment significantly reduced the amount of bioactive IL-1β secreted by BMDMs (Fig 3E). To determine if polarization state altered inflammasome activation by PIM447, BMDMs were skewed toward M1 or M2 prior to inflammasome activation. Interestingly, PIM inhibition significantly decreased inflammasome activation in M2 macrophages (reduced cleaved caspase 1), whereas no signal was detected in M1 macs (Fig. 3D). The first step in activation of the inflammasome is increased transcription of inflammasome proteins by the Nf-κB transcription factor. PIM was previously reported to activate NfκB in cancer cells through direct and indirect mechanisms (31). To determine if PIM activates the inflammasome through NfκB, WT BMDM were treated with PIM447, stimulated with LPS and/or ATP, and the phosphorylation state of NfκB pathway proteins was quantified by densitometry. Treatment with PIM447 decreased IκBα phosphorylation (S32), which corresponded with increased total levels of IκBα, the negative regulator of NfκB. PIM inhibition also decreased phosphorylation of p65 (S536), which is an indicator of p65 transactivation downstream of NFκB (Fig 4F). In addition, PIM447 decreased the mRNA levels of TNFα, an NF-κB target gene (Fig. 4G). Together, these data demonstrate loss of PIM reduces inflammasome activation in macrophages and blocks the release of tumor promoting, pro-inflammatory cytokines.

**Figure 4.**
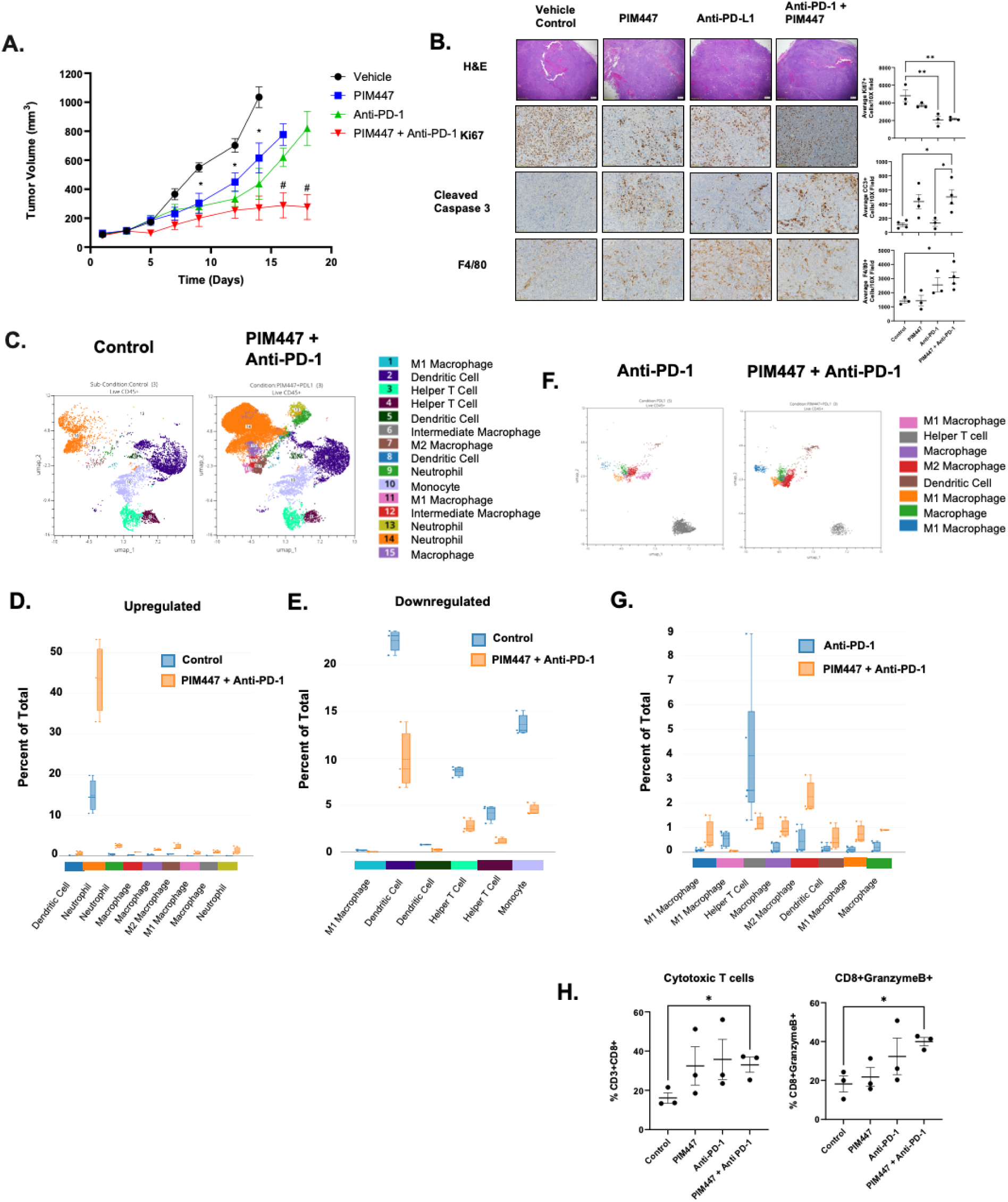
PIM Inhibition Enhances PD-L1 Blockade. A) MyC-Cap tumor-bearing mice (n=8 tumors/group) were treated with vehicle, PIM447, Anti-PD-1, or the combination, and tumor growth was measured over time. B and C) Staining and quantification of Ki67, Cleaved Caspase 3, F4/80, and CD8 in tumors from each cohort. C) UMAP cluster analysis of statistically significant immune cell population changes in control vs PIM447+Anti-PD-1 tumors (n = 3). D) upregulated and E) downregulated immune clusters in PIM447+Anti-PD-1 tumors vs. controls, F and G) UMAP cluster analysis and quantification of immune cell population changes in Anti-PD-1 vs PIM447 + Anti-PD-1 tumors. H) Flow cytometry quantification of CD3+, CD8+, and CD8+GrazymeB+ T cells in each treatment group. (*p<.05,**p<.01).

### PIM inhibition enhances the efficacy of immune checkpoint blockade in a syngeneic model of prostate cancer

Several PIM inhibitors are being actively tested in anti-cancer clinical trials, but none have been tested in combination with anti-PD-1 drugs in prostate cancer. To determine if PIM inhibitors sensitize prostate cancer to immunotherapy, we utilized a syngeneic PCa tumor model. The MyC-Cap prostate cancer cell line was derived from a spontaneous prostate tumor of a c-myc overexpression mouse. MyC-Cap cells have overexpression of wild type androgen receptor (32) as well as androgen receptor splice variant expression (33).Furthermore, MyC-Cap cells are dependent on androgens to grow, however following in vivo androgen withdrawal the cells progress to hormonal independence. MyC-Cap cells are syngeneic and therefore are a useful model to study tumor immunology and immunotherapy. Androgen receptor inhibition has been shown to decrease CD8+ T cell numbers and increase myeloid-derived suppressor cells and PD-L1 expression in murine MyC-Cap prostate tumors (34). PIM1 kinase is co-expressed with c-myc in human prostate cancers and PIM drives c-myc induced prostate cancer progression. Therefore, the MyC-Cap cell line is a particularly relevant cell line to study PIM in an immunocompetent mouse model. One million MyC-Cap PCa cells were injected into the flanks of FVB mice, and tumors were allowed to establish before the mice were divided into the following four treatment groups: vehicle, PIM447 (30 mg/kg, daily, p.o.), anti-PD-1 (200 ug/mo, 3x/wk, i.p.), and PIM447 + anti-PD-1. Treatment with PIM447 or anti-PD-1 alone reduced tumor volume by approximately 20%, whereas combined treatment caused a synergistic anti-tumor response, reducing tumor volume by nearly 80% (Fig. 4A). At the end of the experiment, tumors were isolated from each group for histological examination. Combined treatment with PIM447 and anti-PD-1 significantly decreased proliferation (ki67), increased cell death (cl-caspase 3), and increased overall number of macrophages compared to controls (Fig. 4B). Tumor tissue was also collected from each group for immune profiling using a 15-fluorophore mouse-specific spectral flow cytometry panel (Cytek Biosciences), and FlowSOM, a clustering algorithm, was used to classify our results to produce 30 unique cell clusters. Then, UMAPs were plotted to highlight specific cell cluster changes between control and treatment groups. Treatment with anti-PD-1 alone increased the number of repressive T cells (CD3+CD4+CD8+ T cells) and decreased MDSCs compared to control tumors (Fig. S4A). Treatment with PIM447 alone increased 7 clusters and decreased 3 clusters (Fig. S4B), the majority of which represented macrophage populations. Surprisingly, PIM447 treatment alone increased both M2 (Cdllb+F4/80+Arg1+) and M1 macrophage (Cdllb+F4/80+iNOS+IFNγ+) populations. PIM inhibition also increased the number of neutrophils (Cdllb+Ly6C^lo^Ly6G+iNOS+IFNγ+) and decreased pro-inflammatory monocytes/myeloid-derived suppressor cells (Cdllb+Ly6C+Ly6G+iNOS+IFNγ+). In the combination treatment group, 9 immune cell clusters were upregulated, and 6 clusters were downregulated compared to the control (Fig. 4C). The most significant changes observed in the combination therapy group was a significant increase in anti-tumor neutrophils (Cdllb+Ly6C+Ly6G^int^TNFa^int^IFNg^int^) (Fig. 4D) and a significant reduction in suppressive helper T cells (CD3+CD4+ and CD3+CD4+CD25+) (Fig. 4E). To understand how PIM inhibition primes the TME to favor response to immunotherapy, we isolated cell clusters that significantly changed in the combination group vs Anti-PD-1 alone (Fig. 4F). We observed a significant change in 8 immune cell clusters. Strikingly, nearly all these clusters were macrophage subpopulations (Fig. 4G). Traditional flow cytometry confirmed that PIM inhibition increased the percentage of F4/80+ macrophages, alone and in combination with ICIs (Fig. S4C). Analysis of specific macrophage polarization markers included in our panel indicated that the increased macrophages collected from PIM447 and combination treated tumors are of an intermediate phenotype (Cdllb+F4/80+Arg1+IFNg+ and Cdllb+F4/80+Arg1+TNFα+), suggesting a shift from M2 to M1 macrophages. Importantly, FACS analysis of tumor tissues showed that combination therapy significantly increased the number and activation state (GranzymeB+) of CD8+ T cells compared to all other treatments (Fig. 4H). Collectively, these data indicate that PIM inhibition shifts the immune landscape to favor sensitivity to ICIs by reducing repressive immune cells and increasing cytotoxic T-cell activation.

## Discussion

Growing evidence indicates that PIM kinases have diverse functions in both tumor and immune cells. While PIM is well established as a resistance mechanism in solid tumors, these studies have mainly focused on the pro-survival role of PIM in tumor cells. Here, we utilized next-generation sequencing and whole transcriptomic analysis of tumors isolated from a unique population of treatment-naïve metastatic hormone-sensitive prostate cancer patients to determine how PIM shapes the prostate cancer immune TME and sensitivity to ICIs. The patient population investigated here is valuable because these patients have not received any prior therapy, which avoids inherent changes in the immune TME caused by androgen deprivation therapy and/or chemotherapy. Among the most striking findings from our patient data was that *PIM*-high tumors had significantly higher density of immunosuppressive macrophages compared to *PIM*-low tumors (Fig.1). While most changes were consistent between PIM isoforms in the clustered patient, there were isoform-specific changes. Most notably, *PIM2*-high tumors showed a significant increase in IFNy that was not observed with PIM1 and PIM3-high tumors (Fig S2E). *PIM2*-high tumors were also associated with significantly increased numbers of regulatory T cells and CD4+ T cells, whereas no change was observed in PIM1/3 high samples (Fig S2C). These results suggest that blocking PIM2 may be the most beneficial for improving the anti-tumor response. However, all small molecule PIM inhibitors developed to date are pan-PIM inhibitors, and prior work from our lab and others suggests that blocking all PIM isoforms is needed for a robust anti-tumor response (35). Future studies are warranted to understand how PIM isoforms differentially impact the prostate immune landscape.

PIM kinases are known to support immune cell survival, but their role in macrophage biology and function was unknown. Our results indicate that differential expression of PIM in macrophages can have a significant impact on prostate tumorigenesis. Strikingly, knockout of PIM in macrophages not only blocked the enhanced tumor growth observed in WT co-injected tumors, but actively suppressed tumor growth in an immune competent mouse model (Fig. 2B). Gene expression analysis demonstrated that tumors co-injected with PIM TKO macrophages displayed changes in key anti-tumor signaling activity associated with immunosuppression. Specifically, prostate cancers are known to have low lymphocyte infiltration, which creates a barrier to effective ICI responses. Co-injection of TKO macrophages enhanced pathways related to lymphocyte trafficking, indicating PIM expression in macrophages suppresses adaptive immune responses that contribute to ICI resistance (Fig. 2F). Tumors containing TKO macrophages also showed significantly increased levels of *CCL24*, a prominent chemokine involved in eosinophil recruitment. Increased eosinophils is correlated with improved prognosis in many cancers, including prostate cancer (36). Furthermore, patients with castrate resistant PCa treated with Sipuleucel-T, a cancer vaccine, showed an increase in eosinophil counts, which correlated with improved survival and enhanced T cell responses (37). Thus, eliminating PIM from macrophage shifts the tumor immune microenvironment to favor an anti-tumor response.

Tumor associated macrophages are a predictor of resistance to ICIs in PCa. TAMs are very plastic and can transform in response to stimulation with lipids, cytokines, chemokines, and small bioactive molecules. Genetic or pharmacological loss of PIM activity in BMDM increased the extent of M1 polarization but had no significant effect on M2 signaling pathways. In our *in vivo* studies, treatment with a PIM inhibitor significantly increased the total number of macrophages, with the emergence of intermediate phenotypes with varying expression of both M1 and M2 markers. Our biochemical data indicate that PIM inhibitors skew naïve and M2 TAMs toward an anti-tumor M1 phenotype. Thus, it is likely that the intermediate macrophage phenotypes that emerge after PIM447 treatment are the result of reprogramming of M2 TAMs. However, it is worth noting that PIM is highly involved in many aspects of cytokine signaling. PIM is a transcriptional target of STATs and an upstream activator of suppressor of cytokine signaling (SOCS) proteins, which provides a negative feedback loop to suppress JAK/STAT signaling.(38,39) Therefore, it is possible that the observed increase in M1 and M2 makers observed following PIM inhibitor treatment could be the result of compensatory feedback mechanisms.

Growing evidence indicates that chronic inflammation is a driver of cancer progression and is associated with resistance to ICIs. Our data demonstrate that inhibition of PIM in macrophages significantly reduced inflammatory cytokine production (Figs 2 and 3). Mechanistically, PIM inhibition suppressed activation of the inflammasome and secretion of key cytokines, including IL-1β (Fig. 3D). Biochemical analysis indicate that PIM enhances inflammasome activation by increasing transcriptional activation of NF-kB, which enhances the transcription of canonical inflammasome genes that are associated with inflammation, cell survival and proliferation (31,40,41). NLRP3 inflammasome activation and downstream IL-1β secretion have been associated with PCa progression, but the mechanisms responsible for their dysregulation remains unclear (9). Our results demonstrate that PIM activates macrophage-specific Nf-KB and inflammasome signaling, and treating with PIM inhibitors suppresses chronic inflammation that drives prostate cancer progression (Fig. 3F). These results support recently published data showing that loss of PIM1 can suppress inflammasome activation in BMDMs (42). PIM inhibitors are actively being tested in clinical trials for both hematological and solid tumors. Targeting PIM has shown to enhance the efficacy of chemotherapy and targeted agents (i.e., anti-PI3K, anti-VEGF). Recently, other groups have combined PIM inhibitors with ICIs in pre-clinical studies of melanoma, hepatocellular carcinoma, and triple negative breast cancer (TNBC) (19,43–45). These studies indicated that combination treatment with PIM inhibitor and anti-PD-L1 decreased MDSC and M2 macrophages (19). In our study, immune profiling showed that PIM447 in combination with anti-PD-1 reduced tumor growth by increasing the total number of macrophages and significantly increased CD8+ T cell activity (Fig. 4). Another notable difference in our results is that we observed a robust population of macrophages with an intermediate M1/M2 phenotype following PIM inhibitor treatment. The functional output of these macrophages remains to be determined, but they expressed relatively low levels of M2 markers and high levels of M1 markers, suggesting that inhibition of PIM initiates the transition to an anti-tumor phenotype. Overall, our results support a new role for PIM kinases in regulating TAM polarization and chronic inflammation in the prostate TME. Importantly, we provide preclinical evidence that combining ICIs with PIM inhibitors is an effective strategy to sensitive PCa tumors to immunotherapy.

## Acknowledgement

These studies were supported by funding from the Department of Defense Prostate Cancer Research Program (W81XWH-19-1-0455) and Arizona Biomedical Research Center (ABRC) New Investigator Award (CTR056054) to N.W, and Support the National Cancer Institute (T32CA009213-41A1) to A.C. This work was also supported by the Arizona Health Sciences’ Personalized Defense Initiative and a Cancer Center Support grant from the National Institute of Health (P30CA023074). We would like to thank the following shared resources at the University of Arizona Cancer Center for their help and support: EMSR, TACMASR, FICMSR, and BBSR.

---

**Figure S1.**
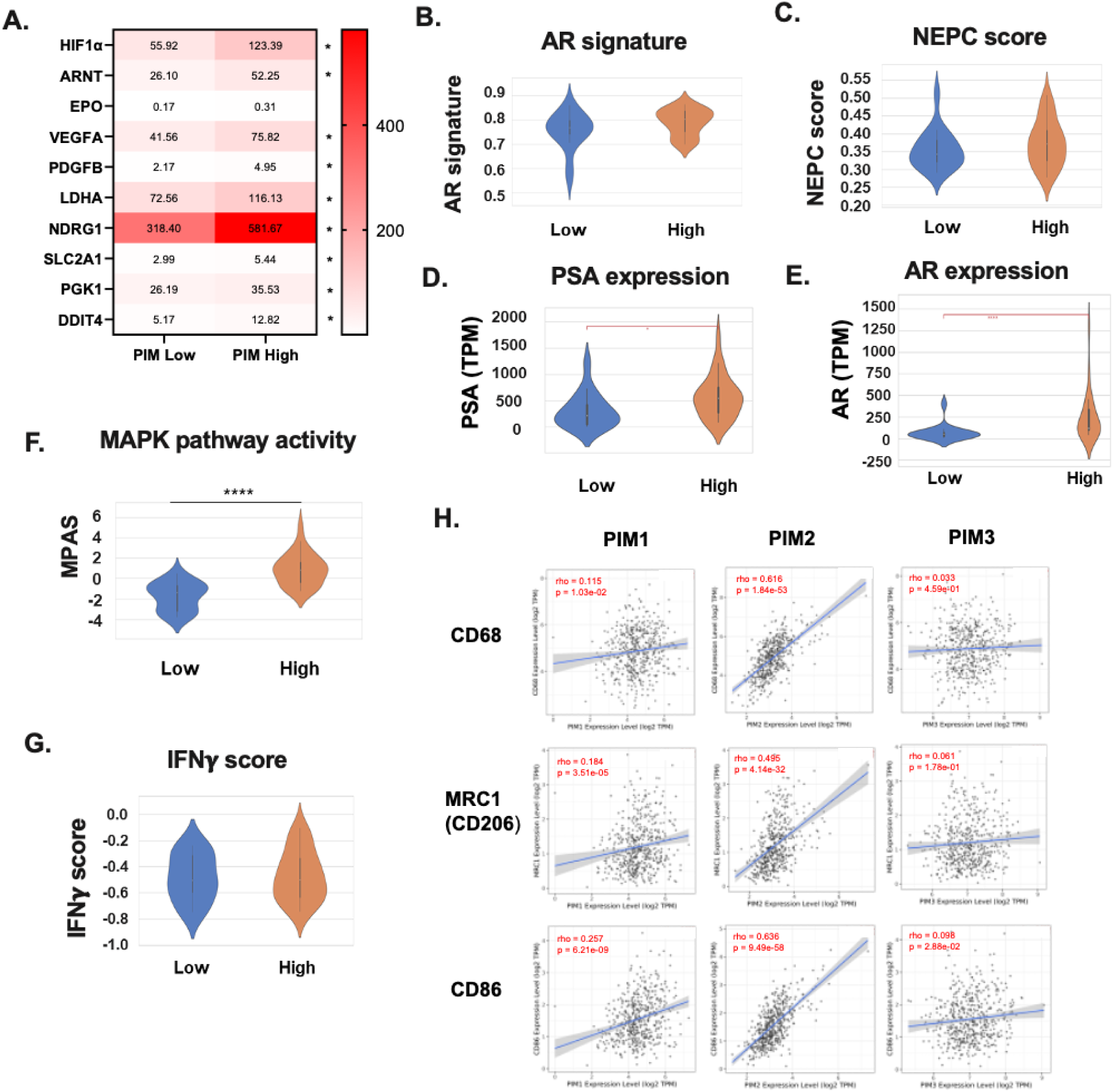
PIM expression correlates with markers of disease progression in human prostate cancer. A) Heatmap of hypoxia-related gene expression in *PIM*-high and *PIM*-low tumors. Comparison onf gene expression in PIM-high vs. PIM–low tumors for: B) Androgen Receptor (AR) signature and C) Neuroendocrine Prostate Cancer (NEPC) score, D) Prostate Specific Antigen (PSA), E) AR, F) MAPK activation score (MPAS), and G) IFNγ score. H) PIM1, PIM2, and PIM3 gene expression was compared to CD68 (pan-macrophage), CD206 (M2 macrophage), and CD86 (M1 Macrophage) using The Cancer Genome Atlas (TCGA) prostate adenocarcinoma (PRAD) data (n=498). ****p<.0001, ***p<.001, **p<.01, *p<.05.

**Figure S2.**
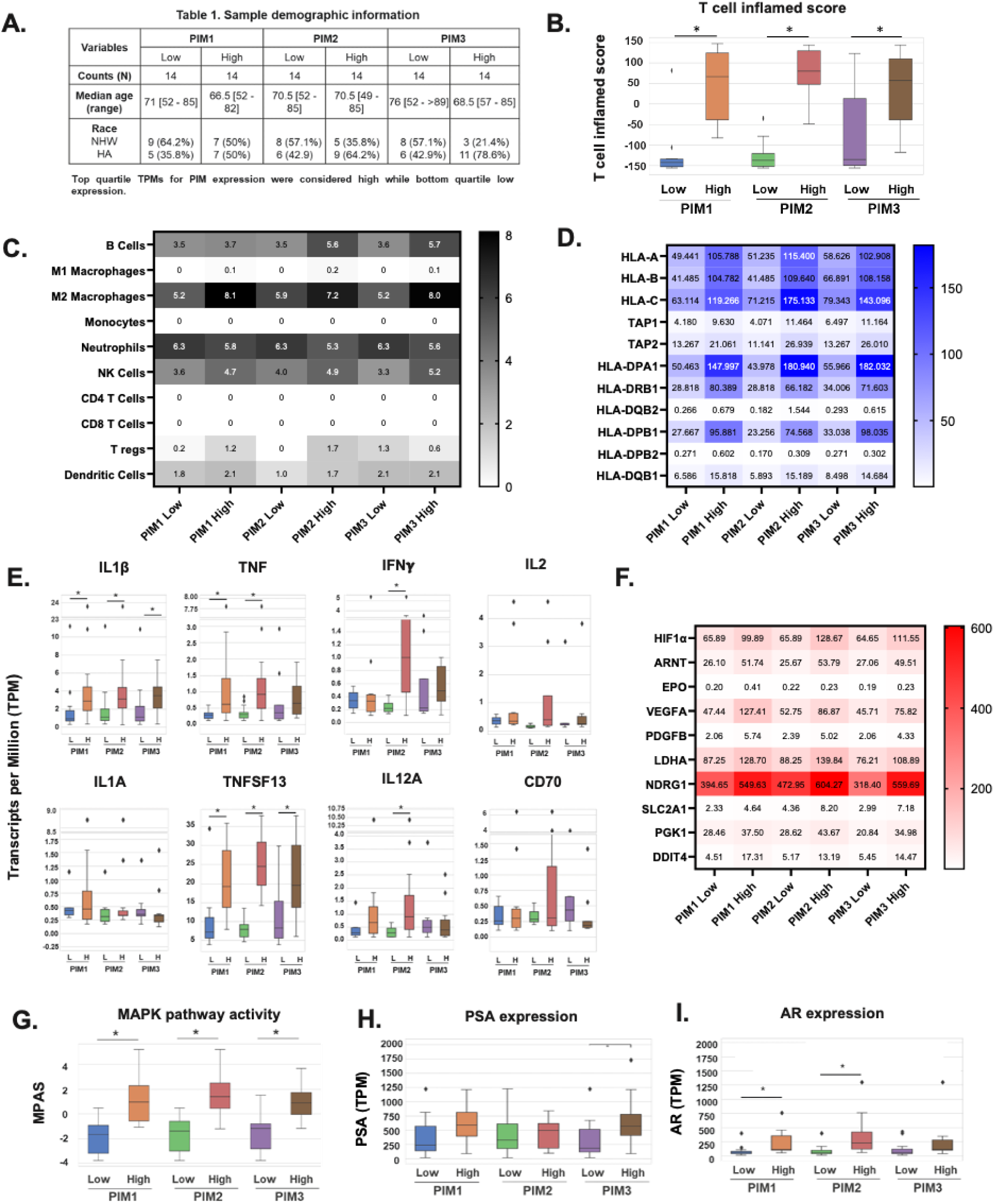
Effect of PIM1/2/3 isoforms on the prostate immune microenvironment. 77 primary prostate and 11 metastatic lymph node samples from treatment-naive metastatic hormone-sensitive Pca patients were analyzed by DNA next-gen seq and WTS. Patient tumros were classified as *PIM1, PIM2, or PIM3*-high(H) and -low(L) by top and bottom quartile. A) Patient demographic information. Comparison of B) T Cell Inflamed Score. C) Heat Maps represents the median % of immune cell fraction of D) MHC Class I and Class II expression, E) Immune gene expression, and F) Hypoxia-gene expression. Comparison of counts for G) MAPK pathway activity, H) Prostate Specific Antigen (PSA), and Androgen Receptor (AR) expression in *PIM1*, *PIM2*, and *PIM3*-high vs low tumors. ****p<.0001, ***p<.001, **p<.01, *p<.05.

**Figure S3.**
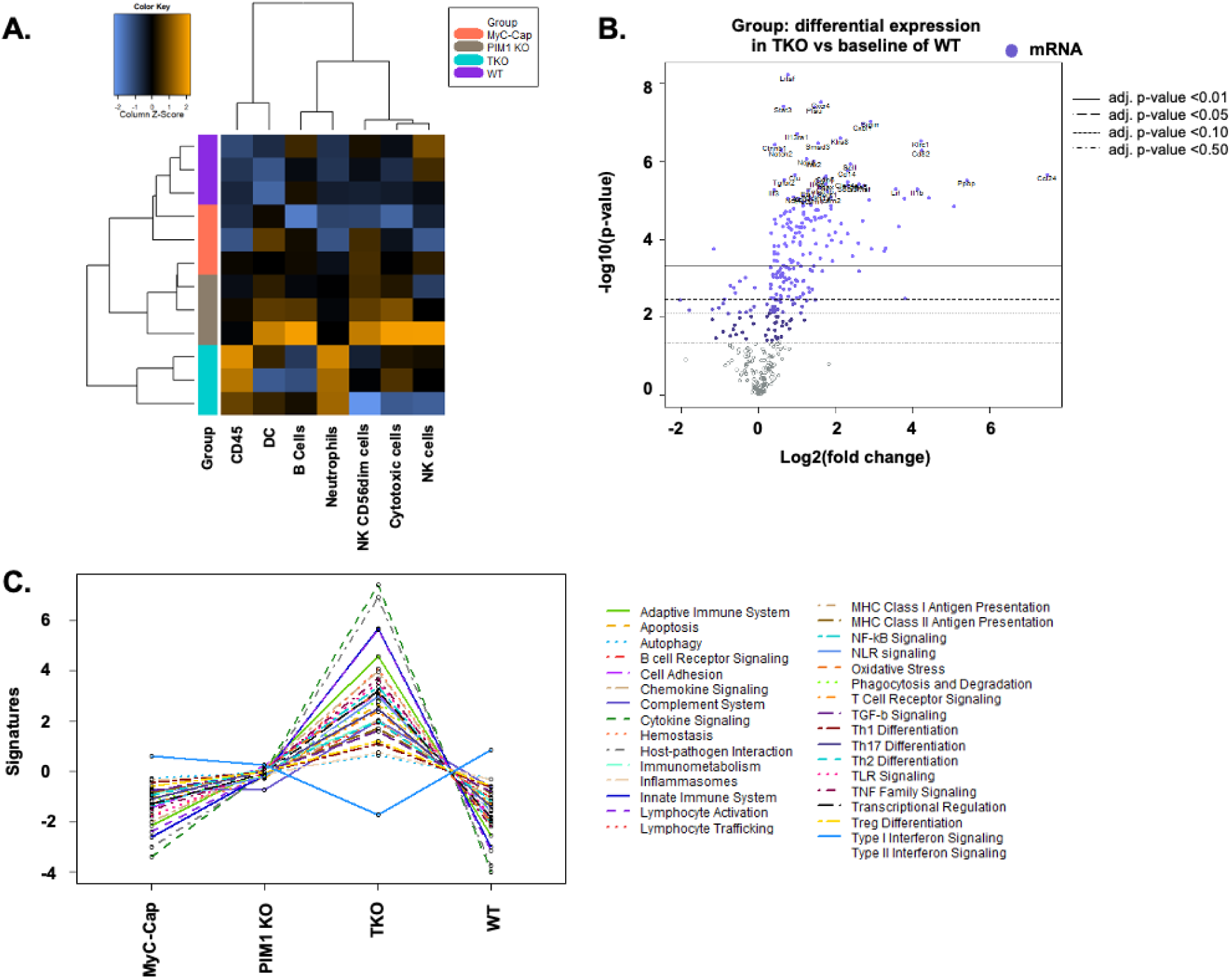
PIM knockout in tumor associated macrophages increases immune cell infiltration, cytokine expression, and immune regulatory signaling pathways. A) Heat map of differentially expressed immune cell populations (Color changes represent Z-score). B) Volcano plot of differential gene expression in TKO vs WT co-injected tumors. C) Pathway signatures from co-injected tumors.

**Figure S4.**
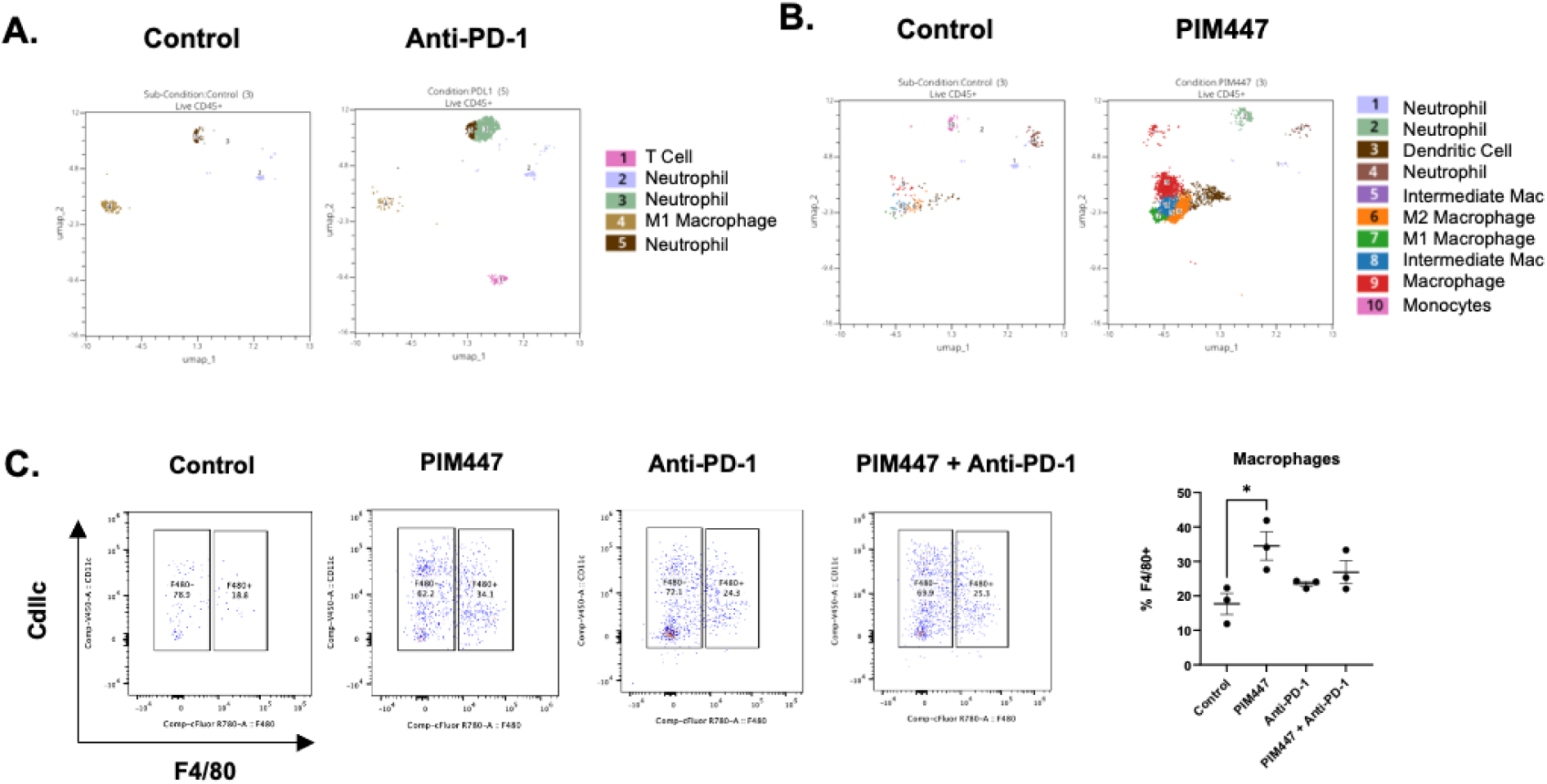
PIM Inhibition Enhances PD-L1 Blockade in a Syngeneic Mouse Model of Prostate Cancer. UMAP analysis of significant changes in A) Control vs Anti-PD-1 and B) Control vs PIM447 treated tumors, C) Representative flow plots and quantification of F4/80+ cells in control and treatment groups.

## References

1. Nonomura N, Takayama H, Nakayama M, Nakai Y, Kawashima A, Mukai M, et al. Infiltration of tumour-associated macrophages in prostate biopsy specimens is predictive of disease progression after hormonal therapy for prostate cancer. BJU Int 2011;107(12):1918–22 doi 10.1111/j.1464-410X.2010.09804.x.

2. Zarif JC, Baena-Del Valle JA, Hicks JL, Heaphy CM, Vidal I, Luo J, et al. Mannose Receptor-positive Macrophage Infiltration Correlates with Prostate Cancer Onset and Metastatic Castration-resistant Disease. Eur Urol Oncol 2019;2(4):429–36 doi 10.1016/j.euo.2018.09.014.

3. DeNardo DG, Ruffell B. Macrophages as regulators of tumour immunity and immunotherapy. Nat Rev Immunol 2019;19(6):369–82 doi 10.1038/s41577-019-0127-6.

4. Martori C, Sanchez-Moral L, Paul T, Pardo JC, Font A, Ruiz de Porras V, et al. Macrophages as a Therapeutic Target in Metastatic Prostate Cancer: A Way to Overcome Immunotherapy Resistance? Cancers (Basel) 2022;14(2) doi 10.3390/cancers14020440.

5. Coschi CH, Juergens RA. Overcoming Resistance Mechanisms to Immune Checkpoint Inhibitors: Leveraging the Anti-Tumor Immune Response. Curr Oncol 2023;31(1):1–23 doi 10.3390/curroncol31010001.

6. Wen Y, Zhu Y, Zhang C, Yang X, Gao Y, Li M, et al. Chronic inflammation, cancer development and immunotherapy. Front Pharmacol 2022;13:1040163 doi 10.3389/fphar.2022.1040163.

7. Abida W, Cheng ML, Armenia J, Middha S, Autio KA, Vargas HA, et al. Analysis of the Prevalence of Microsatellite Instability in Prostate Cancer and Response to Immune Checkpoint Blockade. JAMA Oncol 2019;5(4):471–8 doi 10.1001/jamaoncol.2018.5801.

8. Broz P, Dixit VM. Inflammasomes: mechanism of assembly, regulation and signalling. Nature Reviews Immunology 2016;16(7):407–20 doi 10.1038/nri.2016.58.

9. Xu Z, Wang H, Qin Z, Zhao F, Zhou L, Xu L, et al. NLRP3 inflammasome promoted the malignant progression of prostate cancer via the activation of caspase-1. Cell Death Discov 2021;7(1):399 doi 10.1038/s41420-021-00766-9.

10. Dwivedi S, Goel A, Natu SM, Mandhani A, Khattri S, Pant KK. Diagnostic and prognostic significance of prostate specific antigen and serum interleukin 18 and 10 in patients with locally advanced prostate cancer: a prospective study. Asian Pac J Cancer Prev 2011;12(7):1843–8.

11. Eiró N, Bermudez-Fernandez S, Fernandez-Garcia B, Atienza S, Beridze N, Escaf S, et al. Analysis of the expression of interleukins, interferon β, and nuclear factor-κ B in prostate cancer and their relationship with biochemical recurrence. J Immunother 2014;37(7):366–73 doi 10.1097/cji.0000000000000045.

12. Ferrer FA, Miller LJ, Andrawis RI, Kurtzman SH, Albertsen PC, Laudone VP, et al. Vascular endothelial growth factor (VEGF) expression in human prostate cancer: in situ and in vitro expression of VEGF by human prostate cancer cells. J Urol 1997;157(6):2329–33.

13. Tong Y, Cao Y, Jin T, Huang Z, He Q, Mao M. Role of Interleukin-1 family in bone metastasis of prostate cancer. Front Oncol 2022;12:951167 doi 10.3389/fonc.2022.951167.

14. Wang D, Cheng C, Chen X, Wang J, Liu K, Jing N, et al. IL-1β Is an Androgen-Responsive Target in Macrophages for Immunotherapy of Prostate Cancer. Adv Sci (Weinh) 2023;10(17):e2206889 doi 10.1002/advs.202206889.

15. Chen WW, Chan DC, Donald C, Lilly MB, Kraft AS. Pim family kinases enhance tumor growth of prostate cancer cells. Mol Cancer Res 2005;3(8):443–51 doi 10.1158/1541-7786.MCR-05-0007.

16. Dhanasekaran SM, Barrette TR, Ghosh D, Shah R, Varambally S, Kurachi K, et al. Delineation of prognostic biomarkers in prostate cancer. Nature 2001;412(6849):822-6 doi 10.1038/35090585.

17. Szydłowski M, Prochorec-Sobieszek M, Szumera-Ciećkiewicz A, Derezińska E, Hoser G, Wasilewska D, et al. Expression of PIM kinases in Reed-Sternberg cells fosters immune privilege and tumor cell survival in Hodgkin lymphoma. Blood 2017;130(12):1418–29 doi 10.1182/blood-2017-01-760702.

18. Szydłowski M, Dębek S, Prochorec-Sobieszek M, Szołkowska M, Tomirotti AM, Juszczyński P, et al. PIM Kinases Promote Survival and Immune Escape in Primary Mediastinal Large B-Cell Lymphoma through Modulation of JAK-STAT and NF-κB Activity. Am J Pathol 2021;191(3):567–74 doi 10.1016/j.ajpath.2020.12.001.

19. Xin G, Chen Y, Topchyan P, Kasmani MY, Burns R, Volberding PJ, et al. Targeting PIM1-Mediated Metabolism in Myeloid Suppressor Cells to Treat Cancer. Cancer Immunol Res 2021;9(4):454–69 doi 10.1158/2326-6066.cir-20-0433.

20. Mikkers H, Nawijn M, Allen J, Brouwers C, Verhoeven E, Jonkers J, et al. Mice deficient for all PIM kinases display reduced body size and impaired responses to hematopoietic growth factors. Mol Cell Biol 2004;24(13):6104–15 doi 10.1128/mcb.24.13.6104-6115.2004.

21. Fox CJ, Hammerman PS, Thompson CB. The Pim kinases control rapamycin-resistant T cell survival and activation. J Exp Med 2005;201(2):259–66 doi 10.1084/jem.20042020.

22. van Lohuizen M, Verbeek S, Krimpenfort P, Domen J, Saris C, Radaszkiewicz T, et al. Predisposition to lymphomagenesis in pim-1 transgenic mice: cooperation with c-myc and N-myc in murine leukemia virus-induced tumors. Cell 1989;56(4):673–82 doi 10.1016/0092-8674(89)90589-8.

23. Baytel D, Shalom S, Madgar I, Weissenberg R, Don J. The human Pim-2 proto-oncogene and its testicular expression. Biochim Biophys Acta 1998;1442(2-3):274–85 doi 10.1016/s0167-4781(98)00185-7.

24. Jiménez-García MP, Lucena-Cacace A, Robles-Frías MJ, Ferrer I, Narlik-Grassow M, Blanco-Aparicio C, et al. Inflammation and stem markers association to PIM1/PIM2 kinase-induced tumors in breast and uterus. Oncotarget 2017;8(35):58872–86 doi 10.18632/oncotarget.19438.

25. Jiménez-García MP, Lucena-Cacace A, Robles-Frías MJ, Narlik-Grassow M, Blanco-Aparicio C, Carnero A. The role of PIM1/PIM2 kinases in tumors of the male reproductive system. Sci Rep 2016;6:38079 doi 10.1038/srep38079.

26. Daenthanasanmak A, Wu Y, Iamsawat S, Nguyen HD, Bastian D, Zhang M, et al. PIM-2 protein kinase negatively regulates T cell responses in transplantation and tumor immunity. J Clin Invest 2018;128(7):2787–801 doi 10.1172/jci95407.

27. Casillas AL, Chauhan SS, Toth RK, Sainz AG, Clements AN, Jensen CC, et al. Direct phosphorylation and stabilization of HIF-1α by PIM1 kinase drives angiogenesis in solid tumors. Oncogene 2021;40(32):5142–52 doi 10.1038/s41388-021-01915-1.

28. Liu X, Quan N. Immune Cell Isolation from Mouse Femur Bone Marrow. Bio Protoc 2015;5(20) doi 10.21769/bioprotoc.1631.

29. Korbecki J, Kojder K, Simińska D, Bohatyrewicz R, Gutowska I, Chlubek D, et al. CC Chemokines in a Tumor: A Review of Pro-Cancer and Anti-Cancer Properties of the Ligands of Receptors CCR1, CCR2, CCR3, and CCR4. Int J Mol Sci 2020;21(21) doi 10.3390/ijms21218412.

30. Decker T, Kovarik P. Serine phosphorylation of STATs. Oncogene 2000;19(21):2628–37 doi 10.1038/sj.onc.1203481.

31. Nihira K, Ando Y, Yamaguchi T, Kagami Y, Miki Y, Yoshida K. Pim-1 controls NF-kappaB signalling by stabilizing RelA/p65. Cell Death Differ 2010;17(4):689–98 doi 10.1038/cdd.2009.174.

32. Watson PA, Ellwood-Yen K, King JC, Wongvipat J, Lebeau MM, Sawyers CL. Context-dependent hormone-refractory progression revealed through characterization of a novel murine prostate cancer cell line. Cancer Res 2005;65(24):11565–71 doi 10.1158/0008-5472.Can-05-3441.

33. Ellis L, Ku S, Li Q, Azabdaftari G, Seliski J, Olson B, et al. Generation of a C57BL/6 MYC-Driven Mouse Model and Cell Line of Prostate Cancer. Prostate 2016;76(13):1192–202 doi 10.1002/pros.23206.

34. Xu P, Yang JC, Chen B, Nip C, Van Dyke JE, Zhang X, et al. Androgen receptor blockade resistance with enzalutamide in prostate cancer results in immunosuppressive alterations in the tumor immune microenvironment. J Immunother Cancer 2023;11(5) doi 10.1136/jitc-2022-006581.

35. Torres-Ayuso P, Katerji M, Mehlich D, Lookingbill SA, Sabbasani VR, Liou H, et al. PIM1 targeted degradation prevents the emergence of chemoresistance in prostate cancer. Cell Chem Biol 2024;31(2):326–37.e11 doi 10.1016/j.chembiol.2023.10.023.

36. Luna-Moré S, Florez P, Ayala A, Diaz F, Santos A. Neutral and acid mucins and eosinophil and argyrophil crystalloids in carcinoma and atypical adenomatous hyperplasia of the prostate. Pathol Res Pract 1997;193(4):291–8 doi 10.1016/s0344-0338(97)80006-4.

37. McNeel DG, Gardner TA, Higano CS, Kantoff PW, Small EJ, Wener MH, et al. A transient increase in eosinophils is associated with prolonged survival in men with metastatic castration-resistant prostate cancer who receive sipuleucel-T. Cancer Immunol Res 2014;2(10):988–99 doi 10.1158/2326-6066.Cir-14-0073.

38. Chen XP, Losman JA, Cowan S, Donahue E, Fay S, Vuong BQ, et al. Pim serine/threonine kinases regulate the stability of Socs-1 protein. Proc Natl Acad Sci U S A 2002;99(4):2175–80 doi 10.1073/pnas.042035699.

39. Peltola KJ, Paukku K, Aho TL, Ruuska M, Silvennoinen O, Koskinen PJ. Pim-1 kinase inhibits STAT5-dependent transcription via its interactions with SOCS1 and SOCS3. Blood 2004;103(10):3744–50 doi 10.1182/blood-2003-09-3126.

40. Shen YM, Zhao Y, Zeng Y, Yan L, Chen BL, Leng AM, et al. Inhibition of Pim-1 kinase ameliorates dextran sodium sulfate-induced colitis in mice. Dig Dis Sci 2012;57(7):1822–31 doi 10.1007/s10620-012-2106-7.

41. Staal J, Beyaert R. Inflammation and NF-κB Signaling in Prostate Cancer: Mechanisms and Clinical Implications. Cells 2018;7(9) doi 10.3390/cells7090122.

42. Zhang Z, Xie S, Qian J, Gao F, Jin W, Wang L, et al. Targeting macrophagic PIM-1 alleviates osteoarthritis by inhibiting NLRP3 inflammasome activation via suppressing mitochondrial ROS/Cl(-) efflux signaling pathway. J Transl Med 2023;21(1):452 doi 10.1186/s12967-023-04313-1.

43. Wang JC, Chen DP, Lu SX, Chen JB, Wei Y, Liu XC, et al. PIM2 expression induced by proinflammatory macrophages suppresses immunotherapy efficacy in hepatocellular carcinoma. Cancer Res 2022 doi 10.1158/0008-5472.can-21-3899.

44. Begg LR, Orriols AM, Zannikou M, Yeh C, Vadlamani P, Kanojia D, et al. S100A8/A9 predicts response to PIM kinase and PD-1/PD-L1 inhibition in triple-negative breast cancer mouse models. Commun Med (Lond) 2024;4(1):22 doi 10.1038/s43856-024-00444-8.

45. Clements AN, Warfel NA. Targeting PIM Kinases to Improve the Efficacy of Immunotherapy. Cells 2022;11(22) doi 10.3390/cells11223700.

